# Genomics in the jungle: using portable sequencing as a teaching tool in field courses

**DOI:** 10.1101/581728

**Authors:** Mrinalini Watsa, Gideon A. Erkenswick, Aaron Pomerantz, Stefan Prost

## Abstract

Genetic research is a rapidly evolving field of study that is increasingly being utilized as a tool for wildlife conservation. However, researchers and science educators in remote areas can often find it difficult to access the latest genetic technologies, often due to a combination of high costs, bulky equipment, and lack of infrastructure. Recent technological innovations are resulting in portable, low-cost instruments that enable next-generation sequencing in remote environments, offering new opportunities to generate a more widespread network of trained conservation scientists, particularly within regions of high biodiversity. What is currently lacking are formalized educational efforts to teach participants in biodiverse areas with hands-on training in molecular biology and real-time DNA sequencing techniques. To address this challenge, we report the design and summarized feedback/outcomes of a conservation genetics field course, called ‘Genomics in the Jungle’, that took place at a field research station in the Amazon rainforest of southeastern Peru. The program was established by a small US-based NGO, Field Projects International, and facilitated by a local eco-tourism company in Peru, Inkaterra. We utilized portable sequencing technologies from Oxford Nanopore Technologies, and in-kind support from the manufacturers MiniPCR, MiniOne Systems, Promega, and New England Biolabs. Participants included a mix of non-Peruvian students and local/regional students, some of which had no prior exposure to a genetics laboratory. Overall, we maintain that portable sequencing technology is democratizing scientific research and conservation efforts, and is a major step forward for science educators and conservationists.

## INTRODUCTION

Genetic technology serves a crucial role in ecology and conservation research programs today (Cino & Ettore, 2018; Haig et al., 2015; Hunter, Hoban, Bruford, Segelbacher, & Bernatchez, 2018). As it becomes increasingly affordable and portable, many applications once restricted to a laboratory can now be conducted directly in the field. At the core of such innovation in the field of genomics is the MinION - a 90g, USB powered device with the capability to sequence over four million strands of DNA in a single run (Leggett & Clark, 2017). Beyond portability, the MinION has two additional significant advantages. First, the sequencer and two flowcells start at an initial cost of $1000, and depending on the application, many samples may be multiplexed onto each flowcell, further reducing per sample costs (Pomerantz et al., 2018; Joshua Quick et al., 2017; Srivathsan et al., 2018). Second, the MinION can sequence > 2 mb (Jain et al., 2018; Josh Quick, 2019), compared to up to 50 kb reads by PacBio’s Single Molecule Real-Time (SMRT) sequencing technology. But the greatest advantage to a conservationist is the possibility of conducting in situ sequencing and thus greatly shortening sample storage times, minimizing sample transport and associated costs, local capacity building, and eradicating the need for exporting biological specimens across country borders. Bringing the science to the sample, effectively turning the paradigm onto its head, also opens doors to local scientists and science education.

The MinION has been deployed in a range of field conditions, from real-time sequencing of endangered wildlife in Ecuadorian forests (Pomerantz et al., 2018) to in situ diagnostics during Ebola (Hoenen et al., 2016; Pennisi, 2016) and Zika outbreaks (Faria et al., 2017). The platform has also been utilized as an effective and affordable teaching tool, but only in a classroom or laboratory setting up to now (Zaaijer & Erlich, 2015; Zeng & Martin, 2017). These advances have created an opportunity for a new generation of conservationists who, empowered by portable sequencing technology, will be better equipped to collect and analyze data with fewer resources directly in the field. Excitingly, the application of portable genetic techniques is an opportunity for foreign and local researchers alike. However, empowering both will require changes in how and where science educators focus their efforts. Here, we propose that training the next generation of conservation geneticists should focus on four key areas: The first is gaining experience through living and working in natural environments. Without conducting fieldwork first-hand, a scientist is unlikely to be able to realistically assess the feasibility of a research project, let alone guide it to fruition (Fleischner, Espinoza, & Gerrish, 2017). Second, a field scientist must be well-versed in the collection of diverse biological specimens, incorporating as sources for biological materials as many major taxonomic groups as possible. Third, should fieldwork occur away from one’s home country, a scientist must learn to work with people of different cultures and backgrounds, since successful field research invariably depends on access to their knowledge and collaboration in the use of their field sites. The fourth and final area of training, we believe, should be in molecular laboratory techniques and sequence analysis, given its power to address modern challenges in the conservation of wildlife and natural habitats (Cino & Ettore, 2018; Haig et al., 2015; Hunter et al., 2018). These are the areas of expertise that will maximize the abilities of the next generation of conservationists to be successful in shifting physical and political landscapes.

In 2018, a collaborative effort between a preeminent Peruvian ecotourism company, Inkaterra Hotels, and conservation research organization based in the USA, Field Projects International (FPI), resulted in the installation of the Amazon rainforest’s first field molecular laboratory called the Green Lab. To inaugurate the Green Lab, FPI hosted a field course on applied genetics and genomics as an introduction for foreign and local researchers and stakeholders to the Green Lab’s potential as a research and teaching facility. The course, titled ‘Genomics in the Jungle’, was run by four instructors and was attended by 15 participants from 6 countries around the world. The course prioritized the participation of local and regional researchers through a scholarship program, which would provide direct capacity building and also opportunities for networking with foreign scientists. The program centered around four sequencing experiments involving high-throughput biodiversity assessment using DNA-barcoding, 16S and 18S rRNA metabarcoding of intestinal microbiota, and double-digest restriction site-associated DNA sequencing (ddRAD). These projects were carried out with samples contributed by local researchers and organizations that agreed to publish resulting data with course participants as co-authors, pending the success of each experiment.

Here we report on the planning and execution of this 2-week field course, with input from course participants and instructors alike, to cover all possible perspectives on the implementation and outcomes of this first-of-its-kind event. We evaluate the outcomes of the course in two ways that should be beneficial to other educators. First, we measure how much the actual course deviated from the course plan, taking into account factors that arise in attempting to run a course entirely in a field setting. We report these deviations, as well as the backups we put in place to ensure the success of the program. Second, we list the program outcomes, some which are completed and some which are still in progress, and share summarized participant and instructor feedback as it pertains to those outcomes.

By publishing the course plan, outcomes, and participant feedback, we invite a discussion on the practicality, as well as the costs and benefits, of providing training in applied field genetics under the following conditions: a) teaching at a remote site in the Amazon rainforest; b) requiring no other pre-requisites for participation other than a minimum age of 18; c) restricting the field course to a 2-week period; and d) splitting course participants into teams to highlight different applications of molecular technology.

## METHODS

### Field Site

This field course took place at the Green Lab, which is located at the Inkaterra Guides Field Station (12°31′ S, 69° 2’ W), an hour’s boat ride from the nearest town of Puerto Maldonado in Peru. Surrounding it is previously unlogged forest, protected as a privately operated conservation concession that was once the site of E.O. Wilson’s renowned entomological survey of 1982-83 resulting in the most diverse arboreal ant fauna ever recorded at the time (Wilson, 1987). Laboratory materials were sourced through a mixture of donations and cost-conscious purchasing, with a focus on simplicity, reliability and replaceability (Appendix A). The lab itself is housed in a two-room wooden cabin with a thatched roof, relatively open to the elements, with inbuilt wooden cabinets for storage. Energy requirements are currently met by a combination of solar panels and generator-based power supply, with the intention of switching entirely to solar energy in the near future. At the time of the field course (July 20 - August 5, 2018), power switched from one service to the next four times during the day, during which all AC-powered devices would have to be switched back on. The full capabilities of the lab are listed in Table 1.

**Table 1:** Laboratory capabilities categorized by function, with reference to the number of samples that can be processed at one time.

| Category | Capability | Scale | Equipment | Refrigeration required? | Notes |
| --- | --- | --- | --- | --- | --- |
| Sample Storage | Samples can be stored frozen at -20 C, at 4 C, or at room temperature | Freezer space: 12 cubic feet; Fridge space: 8 cubic feet | Freezer and standalone fridge/freezer combination | NA | Additional shelf space for storage of samples at room temperature is present. |
| DNA Extraction | In-house DNA extraction protocol based on the International Barcode of Life , with per sample extraction cost of ingredients ~\$2.5 | Spin-columns purchased in packs of 250 | Two 24-well centrifuges, two 6-well minicentrifuges, pipettes (all sizes), water bath and thermoblock. | Some buffers require refrigeration, and one requires freezing | Kit-based extractions can be conducted on site, but the in-house methods are preferred due to extremely reduced costs. Spin-columns are a restricting factor on scale and must be imported. |
| DNA quantification | Assay based method that takes ~5 mins to run | One sample at a time, 500 samples per Quantus assay kit | Quantus flouorometer + assays | Assay requires refrigeration for best shelf life, but can withstand 7-10 days at room temperature | This kit is available in Peru but at nearly double the cost as in the US, so we recommend purchasing and carrying it in by hand. |
| PCR | All PCR ingredients and hot-start Taq available, but mixes can be brought in as well | 40 samples at a time | 8 and 16-well PCR machines | All ingredients need to be kept cold | We do not have RT or QPCR capabilities as yet in this lab |
| Gel electrophoresis | Agarose gel electrophoresis, gels made in-house to any specification | 6 x 15 wells at a time | Gel rigs with inbuilt imagers compatible with cell phones | Loading dye alone is kept cool | Imaging is quick and easy, and 1% gels run in as little as 5 minutes on these devices |
| PCR purification | SPRI bead solution, in-house | Extremely affordable, so no limit to use | 1.5 mL and 96-well magnetic racks | SPRI bead solution and stocks are refrigerated | One batch of this solution produces nearly 50 times as much to work with than commercial vendors |
| Gel purification | Promega gel extraction kit | Kits come in packs of 50-250 | Spin-column based kit extraction | None required | Any kit can be used, and SPRI bead cleanups could also substitute here. Kit-based, spin columns are the limiting factor, purchased outside Peru |
| Library prep and Sequencing | MinION-based sequencing | With a second computer, 2 runs can go simultaenously. | Two MinION sequencers, one 96-barcode kit, and one 8GB (soon to be upgraded to 16GB), SSD Mac OS desktop. | All kit ingredients need to be kept frozen, while flowcells must be refrigerated. | Primary restriction is the possibility that you might have a nonfunctional or subpar flowcell, so best to import/carry by hand as close to use as possible |
| Bioinformatics analysis | Online and offline versions of MiniKnow can be operated | NA | Desktop Mac open for unix-based bioinformatics | NA | We recommend bringing personal laptops to conduct bioinformatics analyses in an operating system you are comfortable working in. |

### Participant Eligibility and Diversity

This field course was open to anyone that met the minimum age requirement of 18 years. In order to minimize challenges associated with instructing participants of vastly different experience levels, we intentionally made sure the instructor to participant ratio was high, allowing individualized discussion and instruction as needed. Additionally, all lectures and activities were presented in English only, although additional small group discussions in Spanish did occur, which meant that basic fluency in English was a requirement for participation in this course.

We chose a 2-week course length, based on instructor and participant availability, as well as the costs of accommodation in the field. Shorter durations are conducive to participation by more diverse audiences (for example, professionals without prescribed summer university breaks) and lower course fees, both of which were priorities in this case. Beyond having few enrollment restrictions, and keeping the course duration short, FPI actively worked to ensure diversity among course participants by offering 4 scholarships (one to a Peruvian participant, two to Latin American participants, and one to a participant from a global audience, who turned out to be from the USA). In addition, three staff members were trained as representatives of the field station (IGFS) at no cost to them.

### Course overview

In brief, the course was designed to progress through different levels of organization in three phases (Table 2, see Appendix B for course syllabus). To begin, participants remained in large groups and received exposure to the environment and local wildlife while also completing essential training in molecular wet lab techniques. In the middle of the course, participants were in smaller groups working on one of the four independent projects. As projects neared completion, participants returned to full group sessions for reviewing results and practicing data analysis.

**Table 2:** Course itinerary breakdown by day and activity.

| Phase | Day | Main activity | Lectures | Progress Quiz |
| --- | --- | --- | --- | --- |
| I | 1 | Transport to site; Introductions |  |  |
|  | 2 | Forest navigation; Intro to field laboratory | 1 |  |
|  | 3 | Field activity; Sample collection methodology, solutions and buffer preparation, DNA isolation part I | 1 |  |
|  | 4 | Field activity; Sample collection methodology, Solutions and buffer preparation, DNA isolation part II | 2 |  |
|  | 5 | Visit collaborators at wildlife rescue center; DNA isolation part III |  | Y |
| II | 6 | Independent projects begin in small groups; interspersed field activities and library prep part I | 1 |  |
|  | 7 | Independent library prep; PCR optimization and trouble shooting |  |  |
|  | 8 | Open field activities; library prep part I and part II |  |  |
|  | 9 | Open day / catch-up day | 1 |  |
|  | 10 | Library prep part III and sequencing |  |  |
|  | 11 | Library prep part III and sequencing |  |  |
| III | 12 | Data analysis; presentation preparation |  | Y |
|  | 13 | Project completion; presentation preparation, laboratory clean-up; final presentations and discussion |  |  |
|  | 14 | Laboratory clean-up; exchange contact information; field site departure |  |  |

Overall, course activities were designed to include 15% field work, 20% lectures, 15% bioinformatics, and 50% laboratory work on independent projects. Participants utilized a mixture of hand-written laboratory notebooks and electronic records to keep notes during the course, and took two quizzes to help instructors evaluate progress and understanding of the materials. Lectures occurred in the evening, while lab activities occurred during the day and on some evenings.

### Case Studies

We designed four research projects for teams of participants to execute as described below, and have published all protocols on <u>Protocols.io</u> for reference. Teams included three to four participants and one instructor as a guide.

### Project 1 – ddRAD

Two primate species (*Leontocebus weddelli* and *Saguinus imperator*) were sampled by FPI as a part of a long-term annual mark-recapture program at a neighboring site, Estación Biológica Río Los Amigos (EBLA) that has been operational since 2009 (full protocol in Watsa et al., 2015). In brief, 79 animals from 14 groups were captured in 2018 during June and July, of which 20 animals were selected to be included in this analysis. Femoral vein blood samples from individually tagged animals were stored on Whatman FTA cards as well as in Longmire’s solution (Longmire, Maltbie, & Baker, 1997) at room temperature. We planned to include closely related mother-offspring pairs, fraternal twin-siblings, and some unrelated individuals for a total of 10 individuals per species. Based on the requirements of prior studies run in a laboratory setting on an Illumina-based platform (Thrasher, Butcher, Campagna, Webster, & Lovette, 2018), we were aiming to generate 100ng of DNA from each individual sample. We followed the protocol outlined by Thrasher et al. (2018), with only a few modifications (Watsa, Erkenswick, Pomerantz, & Prost, 2019). Having no access to Blue Pippin facilities, we carried out size selection by gel electrophoresis instead. Furthermore, we modified our primers to contain adaptors to ONT’s barcodes, so that we could multiplex samples using dual indices: primers were designed based on the indices included in Thrasher et al. (2018), as well as ONT’s 96-barcode kit. Finally, not being limited in terms of read length, we planned to choose a longer read for RAD-tag detection if possible. In order to estimate the number of ddRAD tags to sequence, we used the freely available R package, SimRAD (Lepais & Weir, 2014) along with the previously published *Saguinus imperator* genome (GenBank bioproject: PRJNA399417). The full protocol is available on protocols.io (Watsa et al., 2019).

### Project 2 - 16S metagenomics

This study aimed to test a few different variables by analyzing fecal samples targeting the 16S rRNA region using primers 27F and 1429R (Cusco et al., 2017; Klindworth et al., 2013). First, we compared 3 fecal samples (run in duplicate) each from wild vs. captive animals of two primate species: the howler monkey (*Alouatta seniculus*) and the spider monkey (*Ateles chamek*). For wild-sampled emperor tamarins (*Saguinus imperator*), saddleback tamarins (*Leontocebus weddelli*) and night monkeys (*Aotus nigrifrons*), we compared 2 to 3 identical samples extracted using a commercial kit (Qiagen Power Fecal DNA Kit) vs. the in-house DNA extraction protocol based on the Consortium for the Biodiversity of Life (Klindworth et al., 2013). Additionally, we also ran kit-based extractions of fecal samples from wild-sampled primates from EBLA and captive-sampled primates from the Taricaya Rehabilitation Center. In addition, we ran a blank sample to control for possible contamination during extractions and PCRs. Bioinformatic analyses were carried out using ONT’s WIMP software (Juul et al., 2015). For PCR conditions and a full protocol see Protocols.io (Erkenswick, Prost, Watsa, & Pomerantz, 2019).

### Project 3 - 18s metagenomics

This study was similar to the 16S metagenomics study, except utilizing markers that target hypervariable regions of the 18S rRNA gene in eukaryotic organisms (plants, protozoans, metazoans, etc.). Specifically, two sets of primers were used, the forward and reverse primer pair 574f and 1132r (Hugerth et al., 2014), as well as 566f and 1200r (Hadziavdic et al., 2014). PCR amplification protocols followed what was reported in the literature, except for implementing manufacturer recommendations for the use of GoTaq HotStart Polymerase (ProMega) (See Protocols.io). We conducted three experiments: 1) a comparison of gut eukaryotic diversity in captive versus wild primates; 2) a comparison of gut eukaryotic diversity using two different fecal DNA isolation methods (Qiagen power fecal sample kit versus an in-house protocol); 3) a comparison of gut eukaryotic diversity across several Neotropical primate species. In addition, we ran blank samples to control for possible contamination during extractions and PCRs. For this study, captive primate fecal samples of howler, spider, wooly, and saddleback tamarin monkeys came from the Taricaya Wildlife Rehabilitation Center. Wild primate fecal samples of howler, spider, titi, squirrel, capuchin, saki, saddleback tamarin, and emperor tamarin monkeys came from Field Projects International research programs that take place at the Estacion Biologico Rio Los Amigos. A detailed review of the laboratory protocol can be found on <u>protocol.io</u> (Prost, Erkenswick, Watsa, & Pomerantz, 2019).

### Project 4 - DNA barcoding

In this study, we used a compilation of samples of known and unknown taxonomic identities in a multiplexed reaction across a range of markers. For plant samples, we used matk, rBCL, trnh-psBA and a set of long-range rRNA primers (Krehenwinkel et al., 2019; Kress & Erickson, 2012). For mammals we used a cocktail of cytochrome oxidase 1 primers (Kress & Erickson, 2012). For insects, we used the standard primer pair LepF1 LepR1 for cytochrome oxidase (Hebert, Penton, Burns, Janzen, & Hallwachs, 2004). DNA was extracted from each sample, typically from blood (stored on an FTA Whatman elute card or in Longmire’s solution) or a tissue biopsy using in-house extraction protocols based on Kress et al. (Kress & Erickson, 2012). We then attempted to multiplex all successful samples that had PCR amplicons verified via agarose gel electrophoresis onto a single flowcell run, as per the protocol on protocols.io (Pomerantz, Watsa, Prost, & Erkenswick, 2019). We created consensus sequences for the respective barcodes as described in Pomerantz et al. (2018) and Krehenwinkel et al. (2019).

### Division of Labor Among Instructors

Of the four course instructors, all had significant wet laboratory work experience, three had specialized wildlife sampling expertise, two were experienced in MinION-based sequencing, and one instructor was specialized in bioinformatics. Each instructor gave a minimum of one evening lecture and a maximum of three. Some nights had two short lectures led by different instructors (Table 3). For group projects, the ddRAD team and barcoding team had individual instructors as guides, while the third instructor guided the 16S and 18S teams together. The fourth instructor oversaw basic running of the laboratory while project teams worked, ensuring workspace sterility, access to sterile consumables, making aliquots of reagents, and planning future course modules. At the beginning of the course, to limit overcrowding in the field laboratory, two instructors conducted field activities while the other two provided lab training and instruction, splitting the course into two groups. During course phases II and III, participants self-regulated the number of people in the laboratory working at any one time, and equipment conflicts were minimized by using team sign-up sheets.

**Table 3:** A rundown of lectures and their goals in the course.

| **Lectures** | **Aims** |
| --- | --- |
| Introduction to the Madre de Dios, ITA and FPI | An overview of where we were – and why it was important to learn the context before conducting conservation genomics projects. |
| Genetics and wildlife biology | How have the two intersected? What are some cases in which genetic (not genomic) technology have assisted in conservation biology? |
| Genomics and wildlife biology | A history of the development of genomics techniques, and some novel applications to conservation research. |
| An introduction to case studies | A primer to how the samples used in this course were sourced – laying down context for the individual projects. Introductions to DNA barcoding, SNP finding through ddRADSeq, 16S and 18S metagenomics by the respective instructors leading each group |
| Science communication | How can scientists communicate their goals, process and results to a wide range of audiences? |
| Conservation genetics and the illegal wildlife trade | An introduction to illegal trade in primates within Peru. Specific outcomes of recent research that used genetics to assist in a real-life conservation outcome; focus on pangolin wildlife forensics. |
| Real-time results with the MinION | Viewing as runs proceeded, to learn how to interpret the data produced by MinKNOW during a run |
| Bioinformatics of amplicon sequencing on the MinION | Sequence analysis to produce consensus sequences through multiple rounds of polishing for eventual blasting to NCBI. |

### Course Evaluations

All participants provided feedback on the course in three ways: a small set of questions included on the final quiz taken during the course, and an anonymous post-course questionnaire (PCQ) (Appendix C). The final quiz questions asked for on-the-spot feedback about participants’ favorite and least favorite aspect of the course, and one suggestion for improving the course; note, this feedback could not be made anonymous. In the first section of the PCQ, respondents described on a 5-point scale, from Strongly Disagree to Strongly Agree, their opinion on the impact of lectures and assigned readings, how much of a challenge course activities posted, how involved and knowledgeable course instructors were, overall satisfaction, impact on future goals, and if they would recommend the course to others. In the second section, participants evaluate the level of education at which it would be appropriate to take the course as well as the positive, challenge-oriented, or negative words they would use to describe the experience. In the third and final section of the PCQ, they answered open-ended long-form questions on what they would do to improve the course as well as instructor performance. All PCQ responses were entirely anonymized.

### Ethical Note

The Peruvian Ministry of the Environment (SERFOR) granted annual research and collection permits, and the Animal Studies Committees of the University of Missouri-St. Louis (Protocol #733363-5) approved protocols. This study follows the Animal Behaviour Society Guidelines (Rollin & Kessel, 2006) and American Society of Mammalogists’ Guidelines on wild mammals in research (Sikes & Gannon, 2011).

## RESULTS

### Course Participants

There were a total of 15 participants in this field course from 6 countries, with 40% (6) from Latin and South America, including 4 Peruvian participants, and male: female sex ratio of 1.5. Over half (53%, 8) had advanced degrees and the same proportion were professionals in the field of conservation and wildlife biology. Only 20% (3) were current or recent undergraduates (< 1 year from degree). Of the 15, participants had used some laboratory techniques more than once as follows: 73% (11) had used pipettes, 33% (5) had run PCRs, 47% (7) had run gels, and 7% (1) had sequenced DNA. Overall, 80% (12) and 73% (11) had some field research experience or non-genetics laboratory experience, respectively.

### Course Overview

In general, deviations from the course plan were more frequent as the course proceeded. There were no deviations during the very first phase of large group activities involving excursions to the forest or essential lab skills training. During the second phase, minor laboratory setbacks were encountered, as well as the realization that the introduction to bioinformatic techniques should have been planned for earlier in the course as some participants struggled to understand the experimental design of each project without an idea of subsequent analyses.

We experienced one laboratory setback to do with seemingly failing PCR reactions. As participants spent time rerunning failed reactions, an instructor discovered that several of the laboratory’s transilluminators were not bright enough to reveal the gel bands. Although frustrating, no work was lost, and teams were still able to complete their projects due to built-in catch-up days, and eventually the malfunctioning devices were replaced. In another sense, the equipment malfunction provided an important learning opportunity on how to troubleshoot unexpected results.

A second setback pertained to PCR contamination, evidenced by faint bands appearing in PCR negative controls. This was only observed for the 16S and 18S metagenomic projects. As it was not feasible to troubleshoot contamination at the time, participants and instructors discussed how contamination could be addressed bioinformatically through data analysis, and separate sequencing indices were provided to negative control amplifications accordingly. Indeed, this is a standard way to deal with contamination in many research areas (Salter et al., 2014). Also during this phase, instructors decided to cancel one quiz, in order to provide participants with more time to work on independent research projects.

During phase three several notable changes to the course plan took place. First, although it was intended for all project groups to complete library preparation around the same time, this did not occur; the ddRAD team finished first and began sequencing on Day 10, the taxon barcoding team began sequencing on Day 11, and the 16S and 18S teams started sequencing on Day 12. In all cases, sequencing completed overnight, except for the ddRAD project, which ran for a full 36 hours. The staggered schedule was necessary and helpful to allocate shared resources. However it also presented challenges in uniting the entire course for data analysis and preparing presentations. Ultimately, instructors decided to remove the presentation requirement so that project teams could concentrate on creating cleaned-up digital versions of their laboratory notebooks and drafts of introductions and methods sections for papers that might result from the data. Also, two full-group bioinformatics sessions took place during phase III, utilizing taxon barcode sequencing data available as of Day 12. The first session was an introduction and installation session for all software packages that would be used in an analysis pipeline (Python, Albacore, Nanopolish, CANU, and RACON). Software installation took much longer than anticipated, and was not achieved for approximately half of the participants’ computers, due to compatibility errors or the use of many different operating systems, presenting issues that could not be addressed in the field with limited internet access. In the second session, participants were given a subset of the raw data, and tasked with running the entire data pipeline (Appendix B) and blasting consensus sequences against the NCBI database (nBLAST). In lieu of final presentations, we conducted a class discussion on the status of each project and ideas for course publications.

### Summarized Outcomes: Participants

We received feedback from 10 of the 15 course participants on the post-course questionnaire (PCQ). Minimum, maximum and mode values for section 1 of the PCQ evaluated on a 5-point scale are presented in Table 4 below:

**Table 4:** Minimum, maximum and mode values for section 1 of the PCQ evaluated on a 5-point scale: Strongly Disagree, Disagree, Neutral, Agree, and Strongly Agree.

| **Question** | **Mode** | **Minimum** | **Maximum** |
| --- | --- | --- | --- |
| The readings helped me learn the course material | Agree | Agree | Strongly Agree |
| The lectures helped me learn the course material | Strongly Agree | Agree | Strongly Agree |
| The material covered in the course was challenging to me | Agree | Neutral | Strongly Agree |
| The course activities were challenging to me | Neutral | Neutral | Strongly Agree |
| I received enough attention from the instructor(s) | Strongly Agree | Neutral | Strongly Agree |
| The course instructor(s) were sufficiently knowledgeable about the topics taught | Strongly Agree | Strongly Agree | Strongly Agree |
| Overall, I am satisfied at the end of this course in terms of my expectations when I signed up for it | Strongly Agree | Agree | Strongly Agree |
| This course has helped me figure out more about my interests and possible future goals | Strongly Agree | Agree | Strongly Agree |
| I would recommend this course to someone else | Strongly Agree | Agree | Strongly Agree |

**Table 4:** Themes and the frequency with which they were mentioned in open-ended questions from Section 3 of the PCQ (n=10) as well as feedback from the questions included on the final quiz (n=14).

| Theme | Mentions |
| --- | --- |
| **Positive Themes** |  |
| Most memorable moments were consisted of excursions to the forest, river, or oxbow lake, daily night walks, tree-climbing training | 10 |
| Most memorable moment seeing sequence data come out of MinION or loading the sequencer | 7 |
| Have not heard of another course like this being offered | 3 |
| Enjoyed initial training on lab equipment, such as loading gels and using pipettes | 2 |
| Appreciate the opportunity to go from sample collection to sequencing in a portable lab space | 5 |
| Enjoyed project based learning based on real data | 2 |
| Enjoyed the balance between excursions and laboratory work | 2 |
| Feel that they acquired new or improved their skills in molecular lab work | 14 |
| Enjoyed hearing about conservation genetics applications of next generation sequencing from instructors' personal research and ITA's conservation efforts | 4 |
| Felt that course objectives and methods were clear and straightforward | 8 |

| **Things to be Improved** |  |
| --- | --- |
| Expressed that the genetics “vocabulary” was overwhelming | 7 |
| Felt the course was at a level too advanced for their present education background | 2 |
| Felt that the course could have been better organized in terms of assigning participants final projects | 1 |
| Felt that the course was intense or laboratory work repetitive, with days longer than they would have preferred, ultimately suggesting a course extension of a couple of days | 7 |
| Expressed sharing the lab was a particular challenge, both due to space restrictions and navigating team work | 9 |
| Would have liked more time devoted to bioinformatics, including software provided in advance | 8 |
| Would have liked to have had more detailed protocols in advance to prepare better for each day | 1 |
| Would have liked to actually practice collecting samples from wildlife | 2 |

In Section 2 of the PCQ, participants evaluated the appropriateness of taking this course at various points in their education and careers (Fig. 2). They also overwhelmingly voted for positive markers, moderately supported that the course was also challenging, and showed no support for negative markers (Fig. 3). We summarized the responses in section 3 of the PCQ where participants answered a number of open-ended questions, by distilling the most commonly mentioned themes in this feedback (Table 5).

**Figure 1:**
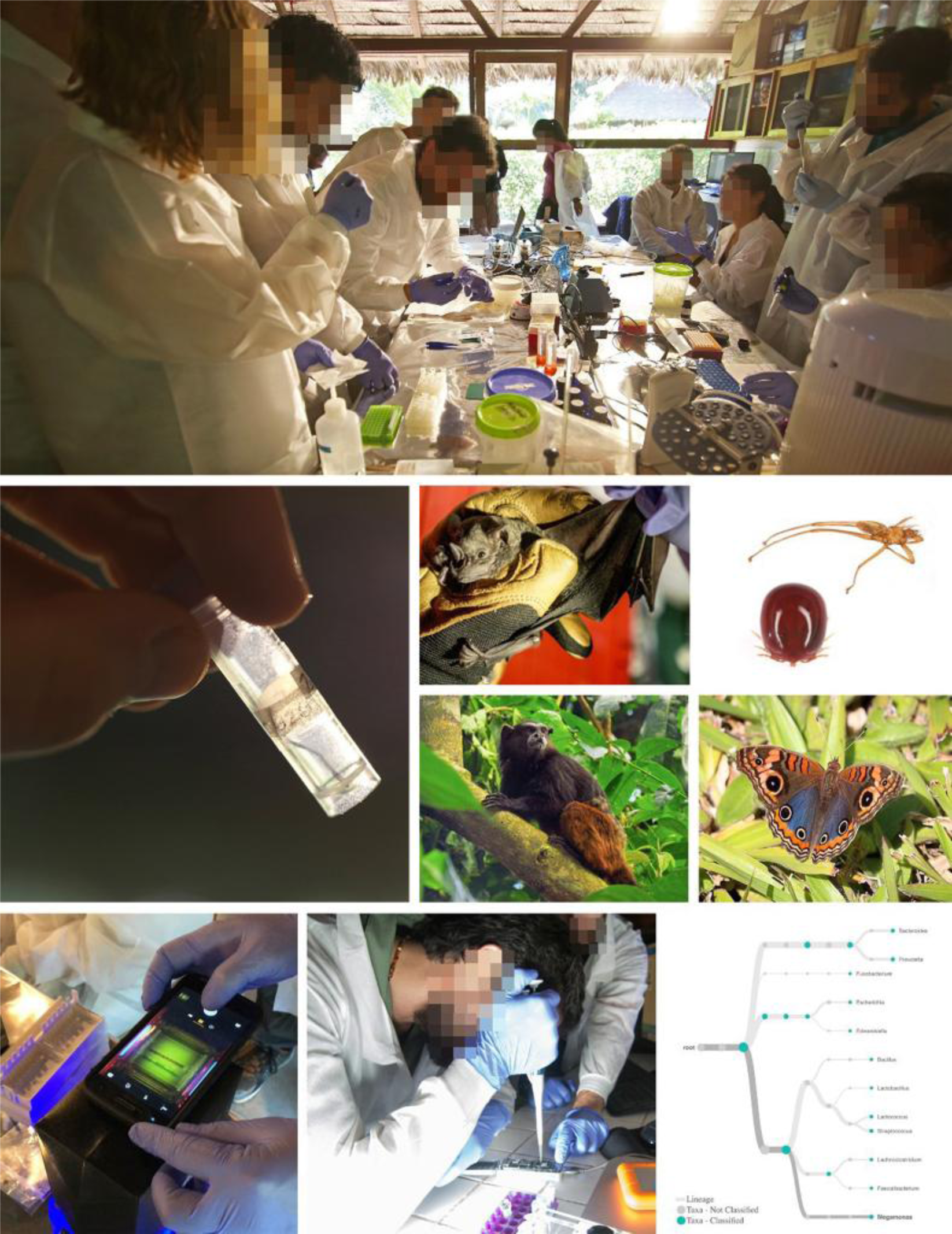
Overview of molecular lab work and example specimens investigated during the field course. Top: Student participants working on independent projects at the Inkaterra Green Lab in southeastern Peru. Middle: example of sample collection and selection of organisms used for genetic analyses during the course, including a bat and its resident arthropod ectoparasites (right), tamarin monkey, and butterfly. Bottom: Student participants running molecular experiments during the course: visualizing PCR products on a portable gel electrophoresis chamber, loading the Oxford Nanopore MinION DNA sequencer, and running a preliminary phylogenetic analysis using data generated from a fecal bacterial sample during the course. Image Credit: Bat: Ryan Peters; Tamarin: Wikimedia creative commons: Gordon E. Robertson; Arthropod ectoparasites, butterfly, and laboratory images: Aaron Pomerantz.

**Figure 2:**
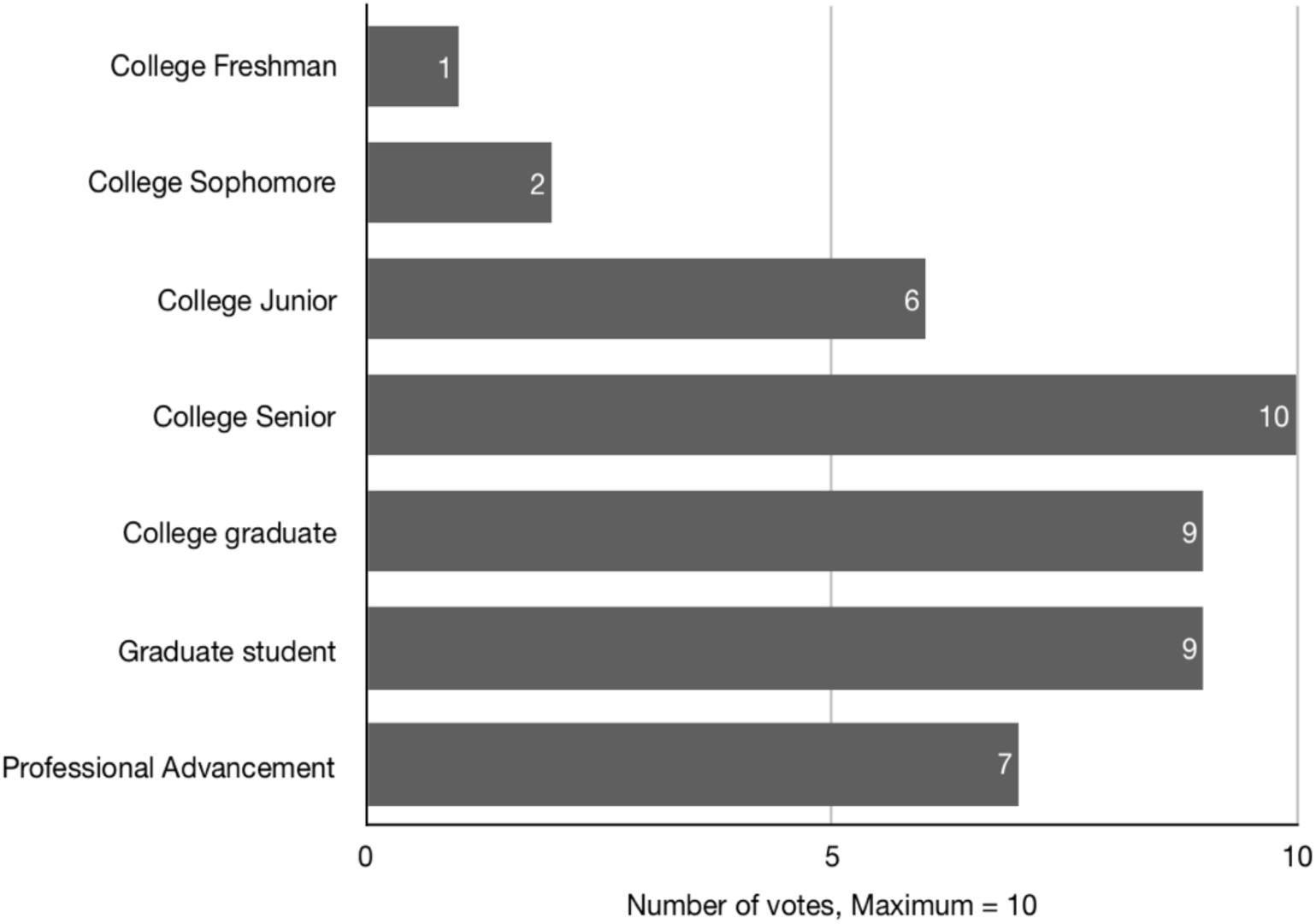
The number of votes received from course participants for each education category at which this course would be appropriate. Maximum survey responses = 10.

**Figure 3:**
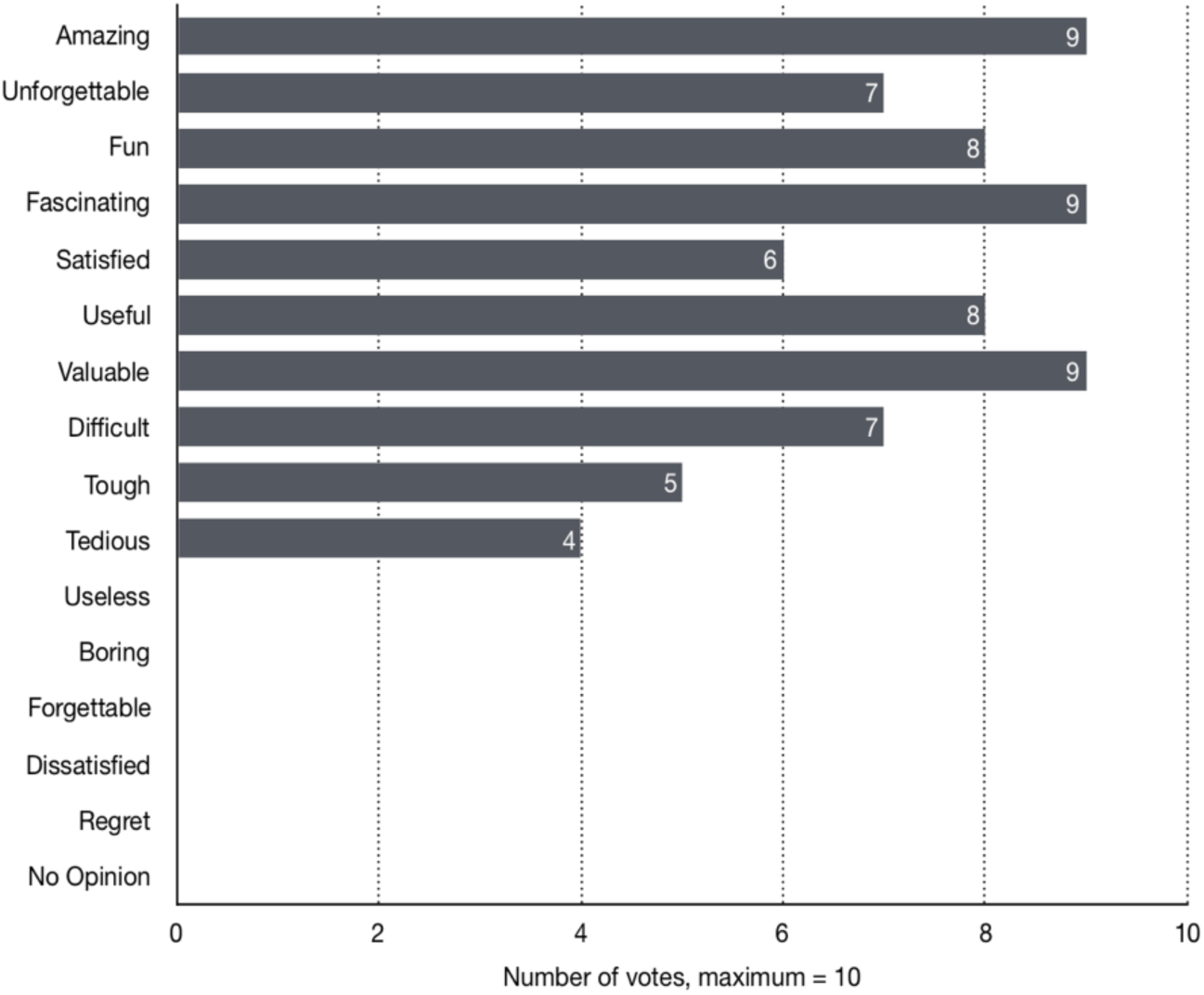
The number of votes received from course participants for positive markers (7), challenge-based markers (3), and negative markers (5) associated with taking this course. Maximum survey responses = 10.

**Figure 4:**
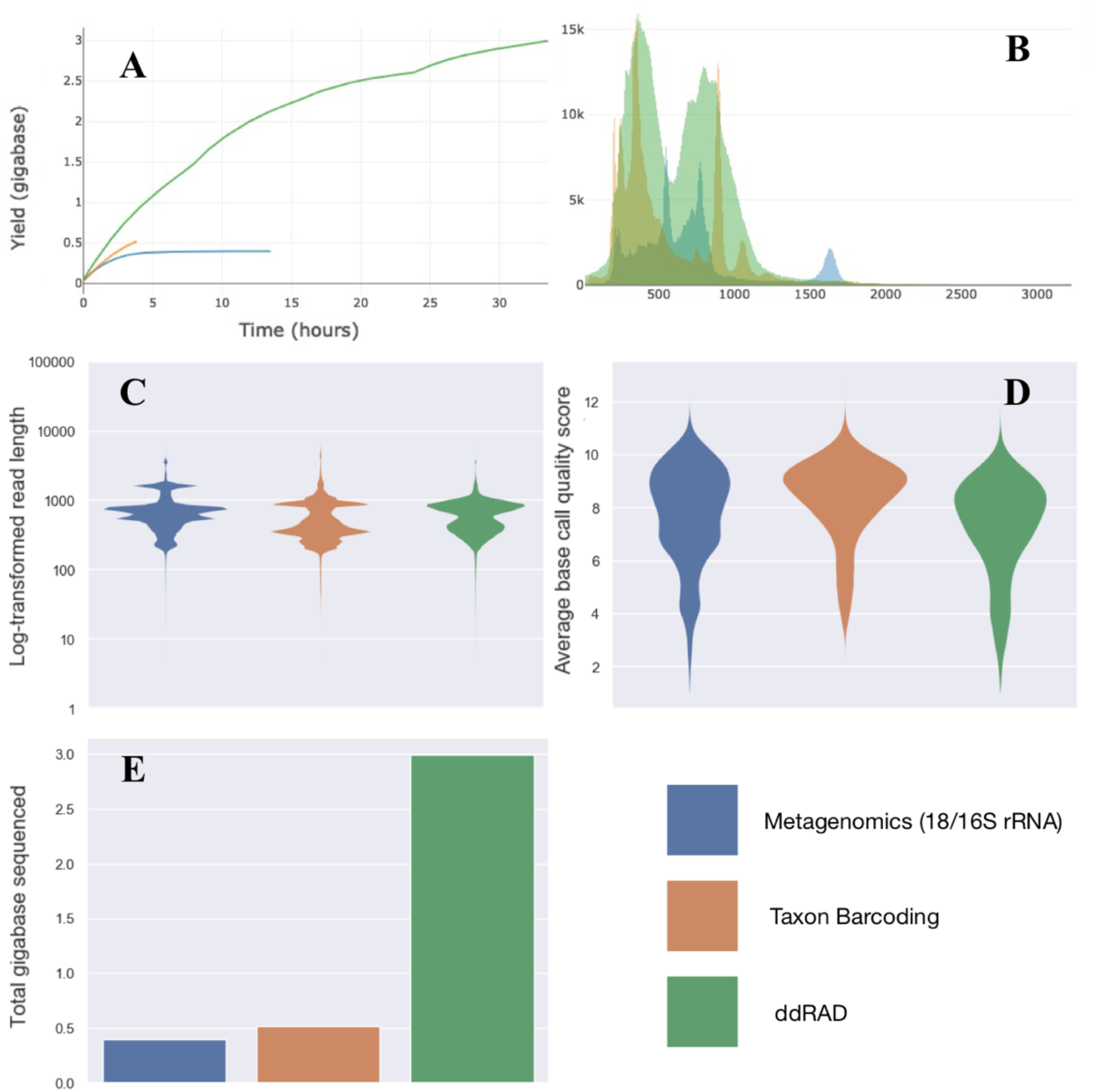
Sequencing performance metrics by project. Plot A: cumulative yields, Plot B: read lengths, Plot C: log-transformed read lengths, Plot D: violin plot of base call quality scores; Plot E: a bar graph of total throughput in gigabases. Plots were generated by NanoComp (De Coster, D’Hert, Schultz, Cruts, & Van Broeckhoven, 2018).

### Summarized Outcomes: Instructors

The four instructors involved in the field course identified key areas of success, as well as, ‘needs for improvement’ with respect to course design and execution. Successes included a diversity of participant backgrounds, the location of the field course, the use of samples collected from regional wildlife to answer legitimate scientific questions, and the use of unstructured working days. They also appreciated having multiples of various laboratory equipment items to allow for uninterrupted workflows by many teams at once.

Obstacles to teaching included the difficulties in comprehensive instruction on bioinformatics within the given timeframe, sharing DNA templates between multiple teams simultaneously, the amount of samples processed, and minimizing sample and reagent contamination with so many people working in an enclosed space that is open to the elements. A proportion of the latter PCRs evidenced contamination of negative controls, which could have been caused by compromised sterility, though some of the reagents may have already possessed small amounts of contamination. Contamination is a known issue among metagenomics studies, even in standard laboratories (Salter et al., 2014). Finally, although it was exciting to run entirely novel experiments (such as the ddRAD project), as a teaching tool it would be more effective to run verified protocols that we know will result in analyzable and publishable data.

### Case Studies

Overall, students and instructors together worked on a series of DNA extractions of samples from feces, blood, and ectoparasites of primates, bats, plants, birds, and insects for a total of 173 extractions. From this pool of extracted DNA, the 16S case study ran 155 PCR reactions, of which they chose 25 amplicon pools for sequencing after running multiple amplifications for the same species. The 18S case study ran 216 PCR reactions and created another 25 amplicon pools over multiple amplifications per species. Both of these teams were able to include comparisons between kit and non-kit extractions, captive and wild primates as well as some inter-species comparisons. The DNA barcoding project ran 192 PCR reactions and multiplexed 112 amplicons onto a flowcell.

We obtained three successful MinION sequencing runs for the four independent research projects conducted during this field course (Table 6): ddRADSeq library (4.6 million reads over 439 active channels in 33 hours, n = 20), 18s and 16s combined (537 thousand reads over 371 active channels in 14 hours n = 50), and DNA barcoding (876 thousand reads over 434 channels in 4 hours, n = 114). Individual publications from each research project are currently in the works, with all course participants welcomed as co-authors.

**Table 6:** Summary statistics of three runs performed during the field course, including mux scan output in each case.

| **Statistic** | **16S_18S_run** | **Barcoding** | **ddRAD** |
| --- | --- | --- | --- |
| Active channels | 371 | 434 | 489 |
| Mean read length | 742.8 | 594.4 | 638.6 |
| Median read length | 657 | 428 | 623 |
| Mean read quality | 7.7 | 8.3 | 7.3 |
| Median read quality | 8.0 | 8.7 | 7.7 |
| Number of reads | 537,631 | 876,109 | 4,692,404 |
| Read length N50 | 779 | 863 | 787 |
| Total bases | 399,356,241 | 520,720,968 | 2,996,466,073 |

## DISCUSSION

Short field courses, in which participants travel to a natural environment to complete a specific training program, can play key roles in guiding career paths, providing local capacity building and drawing attention to a particular environmental problem. In the present case, participants met at a tropical research station in the Peruvian Amazon rainforest to attend a course in applied conservation genetics. The educational backgrounds of the participants ranged from undergraduates to professionals in the field of conservation, even one in the field of engineering, but they all had in common no prior experience carrying out genetic experiences in a remote field site.

### Participants: Course Constraints and Future Recommendations

To everyone’s delight, the vast diversity of participants in terms of their background and prior experience had a number of benefits, and no observed negative impact. Of note among project teams, participants took initiative and pleasure in helping one another as opposed to getting in each other’s way. Most importantly, discussions of career options, conservation initiatives, genomic applications, and science communication, and wildlife biology were frequent, varied and organic in origin.

Both participants and instructors felt that the duration of the course was sufficient for wet laboratory work and lectures, but was not appropriate for practicing bioinformatics to the level that was desired. Dedicating the last two or three days entirely to bioinformatics analyses would be more ideal to accomplish participant software installations, basecalling, exploring run performance, running data pipelines, and interpreting a subset of the results. Reducing the number of case studies could also allow for more time spent on bioinformatics. Furthermore, the use of virtual environments for the bioinformatics pipelines would have mitigated issues arising from participants using different operative systems. Additionally, participants should be encouraged to practice command-line operations prior to arrival.

The readings and lectures enhanced participants’ understanding of the course material, but some reported that new genetics terminology was a challenge. We recommend the creation of a glossary or dictionary for reference that can be provided to incoming participants ahead of time.

### Course Location

That this field course was conducted in the Amazon rainforest’s Green Lab was critical to its success. During travel to and from the Green Lab and through discussion with instructors and citizens of Peru, each participant learned first-hand about regional factors that affect the conservation of biodiversity. Southeastern Madre de Dios faces acute threats from artisanal and small-scale gold mining that include habitat destruction and the bioaccumulation of mercury in local wildlife (Kumar, Divoll, Ganguli, Trama, & Lamborg, 2018; Moreno-Brush et al., 2017). At this time, no comprehensive reference library of DNA barcodes for the fauna and flora of the region exists, although this would greatly enhance biodiversity assessments and monitoring in the face of these threats.

On this field course, a visit to the Taricaya Ecoreserve, a local wildlife research and rehabilitation center, as well as immersion in the conservation efforts of the Inkaterra Association (ITA) provided context for the importance of conservation technology to the local community. Additionally, holding the course in this location created an opportunity for four local researchers to be trained in applied molecular research methodologies and enlarged their professional networks, which we believe will have enduring benefits for the area. Puerto Maldonado, the capital of the Madre de Dios department, is the nearest urban area to the Green Lab (one hour by boat), and is also the location of the Universidad Nacional Amazónica de Madre de Dios. There are currently no genetics training programs available, but the completion of this course has provided a blueprint towards regularly offering certificate training workshops to students in the veterinarian and forest engineering programs.

The field site itself, which had 18 h of electricity, reliable internet access for most of the day, and all facilities located within one hectare of each other, made it possible to seamlessly shift between lectures, laboratory work, personal spaces, and the forest trail system. For example, participants would come off the trail and directly into the laboratory to complete steps in their protocols, and then leave again for a meal, shower, night hike or tree climbing session. Case studies centered on scientific questions to which the answers remained unknown, and utilized biological specimens collected from regional wildlife within the last two years. As such, every project was grounded in reality, with a direct conservation or research impact, providing a clear incentive to success. Consequently, grades and quizzes were not the quintessential motivating factors that they normally are on a course. Instead, the course brought in an additional incentive to take each project eventually to a meaningful finding and publication. With the understanding that returning to the Green Lab for further analyses was not a possibility, the focus was on consolidating all possible information required to analyze and eventually publish project outcomes. This motivated participants to keep clear laboratory records, to write drafts of methods and results sections, and to maintain a high level of integrity in their laboratory work. While this did increase the pressure on the groups to complete projects, unstructured days allowed them the freedom to determine an acceptable balance of work and “play.”

### Field Laboratory Equipment

Initially, participants extracted DNA for all projects together, which went well. Subsequently, however, project teams struggled to keep track of templates which were needed repeatedly by multiple groups. We suggest separate sample streams dedicated to each team, as well as separate reagent stocks to minimize contamination along the way. Contamination of samples did occur, as detected by faint amplification in PCR negative controls, but this also provided an opportunity to discuss how to deal with contamination during data analysis. Having a large group operate in a relatively small laboratory space did cause a strain on the participants at times, as between the project groups, all eight PCRs, five gel rigs, four centrifuges, and five sets of pipettes were in constant use. While there was no appreciable halt in productivity when items malfunctioned due to the number of backups available, PCR rigs were a limiting factor and required signup sheets for fair allocation across teams. Having many 8 and 16-well PCR machines allowed access to all teams simultaneously, however, one additional large-scale machine would alleviate some pressure for optimization and replicate PCRs. The reduction of the number of projects would have certainly alleviated some of the challenges related to laboratory ergonomics and time management.

Two of the instructors on this course pioneered the use of the MinION and an entirely portable laboratory in an Ecuadorian rainforest in 2016 (Pomerantz et al., 2018). In comparison to this scenario, our field course had much more infrastructural support, with on-site refrigeration and larger-scale centrifuges. In order to accommodate all four sequencing projects, we had three MinION sequencers on site, ultimately only using two simultaneously. Since the sequencing might last for up to 72 hours for larger projects, having multiple devices is critical to remaining on schedule.

### Portable Sequencing in the Amazon rainforest

Loading the sequencer and observing the first reads being basecalled live during the run was overwhelmingly one of the most memorable moments in this course. The excitement during these sessions was palpable, and due to the ability to run NCBI BLAST searches online, we were immediately able to verify many of the sequences produced by the DNA barcoding team. Data from the other teams required more analysis steps and was harder to interpret beyond basic run performance parameters. Thus, designing projects with more of an immediate impact in terms of results, based more on amplicon sequencing, can enhance the experience of going from sample to interpreted sequencing results within a two-week timeframe. Downloading databases to BLAST offline would also be advantageous to teach bioinformatics post-experiment.

We experienced a fair amount of variation in run success from project to project, with the best outcome from the ddRAD team who obtained 4.5 million reads in ∼48 hours. Although we had a MacMini with 8GB RAM, a 250GB SSD, and a 2.6 GHz Intel Core i5 processor, the attempted run for the DNA barcoding team failed on this device 1 hour into the run. We were able to simply unplug the MinION and transfer it to a faster laptop to complete the run. The fewest reads (state reads) were unfortunately obtained from the flowcell running the 16s and 18s metagenomics libraries. However, subsequent pilot analyses on WIMP (Juul et al., 2015) indicated some success, but coverage overlap is yet to be assessed.

Ultimately, we strongly recommend utilizing flowcells purchased as close to the date of the course as possible to avoid low read numbers. The newest chemistry (LSK SQ 109) from ONT has resulted in a flowcell that can remain unopened at room temperature for up to 4 weeks, and is covered by a 12 week warranty. Oxford Nanopore Technology has announced a lyophilized field sequencing kit for gDNA (SQK-LRK001) which would allow for transport by hand of the sequencing kit and flowcells without the constraints posed by refrigeration. They have also released information of a MinIT (ONT’s computing device) or the soon-to-be-released MK1c (a combined MinION sequencer, MinIT computer, and screen) that may be able to remove some of the computational challenges of sequencing in remote locations. Even without access to computing servers, ONT’s newly public base-calling software (Guppy; available from https://nanoporetech.com/community) can live-basecall nearly 50% faster than the previous ONT basecaller Albacore 2, allowing for quicker local basecalling of the field course data on dual- or quad-core laptops.

## CONCLUSION

Overall, we maintain that portable sequencing technology and the use of open-source analytical software is democratizing scientific research and conservation efforts, and a major step forward for science educators and conservationists. While these types of courses can be held in any field setting, we believe that their greatest impact will be in places where biodiversity is highest and adequate training of local scientists is lacking. By sharing the feedback and outcomes of this program, a first of its kind in the Amazon, we hope to encourage educators and conservation organizations to consider how the application of modern research and conservation techniques could become much more diffuse than it currently is, with larger proportions of researchers and conservationists that reside in less developed areas doing cutting-edge science despite reduced funding and infrastructure, overall. Our attempt at achieving this also enriched the learning experience for students from more privileged educational systems. Hence, making a concerted effort to teach conservation genetics in the field can be a win-win situation for all students involved, and will likely hasten the implementation of applied conservation genetics. We believe that there is enormous potential to alter the primary way in which conservation genetics has been taking place until now – primarily by foreign groups that transport samples to external laboratories for analysis. In-the-field genetic courses show the potential to foster more localized, on the ground, yet technically modern, conservation genetics work.

## ACKNOWLEDGEMENTS

The authors would like to thank the primary organizations involved in facilitating this program, Field Projects International and the Inkaterra Asociación. We wish to acknowledge the following course participants who gave additional feedback that was used in assessing the field course for this publication: Jan Brack, Si Cave, Dr. Jenny Chen, Samantha Lopez Clinton, David Dainko, Frances Humes, Dr. Yaduraj Khadpekar, Mary McElroy, Dong Jin Park, Mariana Paz, Dr. Nicholas W. Pilfold, Daniel Rudin, Adolfo Schmitt, and Andres Viñas. Additional thanks to the Taricaya Ecoreserve for allowing the course participants to visit the rehabilitation and rescue center, and work with samples from its captive animal population. Finally, we are especially grateful to the following corporations that gave in-kind support to this field course through equipment or reagent donations including Promega, MiniPCR, MiniOne Systems, and New England Biolabs.

## REFERENCES

Cino, P., & Ettore, R. (2018). The ongoing transition at an exponential speed from conservation genetics to conservation genomics. Genetics and Biodiversity Journal, 2(2), 47–54.

Cusco, A., Vines, J., D’Andreano, S., Riva, F., Casellas, J., Sanchez, A., & Francino, O. (2017). Using MinION to characterize dog skin microbiota through full-length 16S rRNA gene sequencing approach. Biorxiv. http://doi.org/10.1101/167015

De Coster, W., D’Hert, S., Schultz, D. T., Cruts, M., & Van Broeckhoven, C. (2018). NanoPack: visualizing and processing long-read sequencing data. Bioinformatics, 34(15), 2666–2669. http://doi.org/10.1093/bioinformatics/bty149

Erkenswick, G. A., Prost, S., Watsa, M., & Pomerantz, A. (2019, March 11). 16S metagenomics in a field setting. Protocols.Io. http://doi.org/10.17504/protocols.io.y2ffybn

Faria, N. R., Quick, J., Claro, I. M., Thézé, J., de Jesus, J. G., Giovanetti, M., et al. (2017). Establishment and cryptic transmission of Zika virus in Brazil and the Americas. Nature Publishing Group, 546(7658), 406–410. http://doi.org/10.1038/nature22401

Fleischner, T. L., Espinoza, R. E., & Gerrish, G. A. (2017). Teaching biology in the field: importance, challenges, and solutions. BioScience, 67(6), 558–567.

Hadziavdic, K., Lekang, K., Lanzen, A., Jonassen, I., Thompson, E. M., & Troedsson, C. (2014). Characterization of the 18S rRNA gene for designing universal eukaryote specific primers. PLoS One, 9(2), e87624–10. http://doi.org/10.1371/journal.pone.0087624

Haig, S. M., Miller, M. P., Bellinger, R., Draheim, H. M., Mercer, D. M., & Mullins, T. D. (2015). The conservation genetics juggling act: integrating genetics and ecology, science and policy. Evolutionary Applications, 9(1), 181–195. http://doi.org/10.1111/eva.12337

Hebert, P. D. N., Penton, E. H., Burns, J. M., Janzen, D. H., & Hallwachs, W. (2004). Ten species in one: DNA barcoding reveals cryptic species in the neotropical skipper butterfly Astraptes fulgerator. Proceedings of the National Academy of Sciences of the United States of America, 101(41), 14812–14817. http://doi.org/10.1073/pnas.0406166101

Hoenen, T., Groseth, A., Rosenke, K., Fischer, R. J., Hoenen, A., Judson, S. D., et al. (2016). Nanopore sequencing as a rapidly deployable Ebola outbreak tool. Emerging Infectious Diseases, 22(2), 331–334. http://doi.org/10.3201/eid2202.151796

Hugerth, L. W., Muller, E. E. L., Hu, Y. O. O., Lebrun, L. A. M., Roume, H., Lundin, D., et al. (2014). Systematic design of 18S rRNA gene primers for determining eukaryotic diversity in microbial consortia. PLoS One, 9(4), e95567. http://doi.org/10.1371/journal.pone.0095567

Hunter, M. E., Hoban, S. M., Bruford, M. W., Segelbacher, G., & Bernatchez, L. (2018). Next-generation conservation genetics and biodiversity monitoring. Evolutionary Applications, 11(7), 1029–1034. http://doi.org/10.1111/eva.12661

Jain, M., Koren, S., Miga, K. H., Quick, J., Rand, A. C., Sasani, T. A., et al. (2018). Nanopore sequencing and assembly of a human genome with ultra-long reads. Nature Biotechnology, 36(4), 338–345. http://doi.org/10.1038/nbt.4060

Juul, S., Izquierdo, F., Hurst, A., Dai, X., Wright, A., Kulesha, E., et al. (2015). What’s in my pot? Real-time species identification on the MinION^TM^. Biorxiv, 030742. http://doi.org/10.1101/030742

Klindworth, A., Pruesse, E., Schweer, T., Peplies, J., Quast, C., Horn, M., & Glockner, F. O. (2013). Evaluation of general 16S ribosomal RNA gene PCR primers for classical and next-generation sequencing-based diversity studies. BioScience, 41(1), 1–11. http://doi.org/10.1093/nar/gks808

Krehenwinkel, H., Pomerantz, A., Henderson, J. B., Kennedy, S. R., Lim, J. Y., Swamy, V., et al. (2019). Nanopore sequencing of long ribosomal DNA amplicons enables portable and simple biodiversity assessments with high phylogenetic resolution across broad taxonomic scale. GigaScience. http://doi.org/10.1093/gigascience/giz006

Kress, W. J., & Erickson, D. L. (2012). DNA Barcodes. (W. J. Kress & D. L. Erickson, Eds.). Humana Press.

Kumar, A., Divoll, T. J., Ganguli, P. M., Trama, F. A., & Lamborg, C. H. (2018). Presence of artisanal gold mining predicts mercury bioaccumulation in five genera of bats (Chiroptera). Environmental Pollution, 236, 862–870. http://doi.org/10.1016/j.envpol.2018.01.109

Leggett, R. M., & Clark, M. D. (2017). A world of opportunities with nanopore sequencing. Journal of Experimental Botany, 68(20), 5419–5429. http://doi.org/10.1093/jxb/erx289

Lepais, O., & Weir, J. T. (2014). SimRAD: an R package for simulation-based prediction of the number of loci expected in RADseq and similar genotyping by sequencing approaches. Molecular Ecology Resources, 14(6), 1314–1321. http://doi.org/10.1111/1755-0998.12273

Longmire, J. L., Maltbie, M., & Baker, R. J. (1997). Use of “lysis buffer” in DNA isolation and its implication for museum collections. Occasional Papers, Museum of Texas Tech University, (163), 1–3.

Moreno-Brush, M., Portillo, A., Brändel, S. D., Storch, I., Tschapka, M., & Biester, H. (2017). Mercury concentrations in bats (Chiroptera) from a gold mining area in the Peruvian Amazon. Ecotoxicology (London, England), 48, 653. http://doi.org/10.1007/s10646-017-1869-1

Pennisi, E. (2016). Pocket DNA sequencers make real-time diagnostics a reality. Science, 351(6275), 800–801. http://doi.org/10.1126/science.351.6275.800

Pomerantz, A., Penafiel, N., Arteaga, A., Bustamante, L., Pichardo, F., Coloma, L. A., et al. (2018). Real-time DNA barcoding in a rainforest using nanopore sequencing: opportunities for rapid biodiversity assessments and local capacity building. GigaScience, 7(4), 74–14. http://doi.org/10.1093/gigascience/giy033

Pomerantz, A., Watsa, M., Prost, S., & Erkenswick, G. A. (2019, March 11). DNA Barcoding in a field setting. Protocols.Io. http://doi.org/10.17504/protocols.io.y2sfyee

Prost, S., Erkenswick, G. A., Watsa, M., & Pomerantz, A. (2019, March 11). 18S metagenomics in a field setting. Protocols.Io. http://doi.org/10.17504/protocols.io.y2ifyce

Quick, Josh. (2019, January 15). Ultra-long read sequencing protocol for RAD004. Protocols.Io. http://doi.org/10.17504/protocols.io.mrxc57n

Quick, Joshua, Grubaugh, N. D., Pullan, S. T., Claro, I. M., Smith, A. D., Gangavarapu, K., et al. (2017). Multiplex PCR method for MinION and Illumina sequencing of Zika and other virus genomes directly from clinical samples. Nature Protocols, 12(6), 1261–1276. http://doi.org/10.1038/nprot.2017.066

Rollin, B. E., & Kessel, M. L. (2006). Guidelines for the treatment of animals in behavioural research and teaching. Animal Behaviour, 71, 245–253. http://doi.org/10.1016/j.anbehav.2005.10.001

Salter, S. J., Cox, M. J., Turek, E. M., Calus, S. T., Cookson, W. O., Moffatt, M. F., et al. (2014). Reagent and laboratory contamination can critically impact sequencebased microbiome analyses. BMC Biology, 12(1), 118–12. http://doi.org/10.1186/s12915-014-0087-z

Sikes, R. S., & Gannon, W. L. (2011). Guidelines of the American Society of Mammalogists for the use of wild mammals in research. Journal of Mammalogy, 92(1), 235–253. http://doi.org/10.1644/10-MAMM-F-355.1

Srivathsan, A., Baloĝlu, B., Wang, W., Tan, W. X., Bertrand, D., Ng, A. H. Q., et al. (2018). A MinION-based pipeline for fast and cost-effective DNA barcoding. Biorxiv, 1–36. http://doi.org/10.1101/253625

Thrasher, D. J., Butcher, B. G., Campagna, L., Webster, M. S., & Lovette, I. J. (2018). Double-digest RAD sequencing outperforms microsatellite loci at assigning paternity and estimating relatedness: A proof of concept in a highly promiscuous bird. Molecular Ecology Resources, 215, 403–13. http://doi.org/10.1111/1755-0998.12771

Watsa, M., Erkenswick, G. A., Halloran, D., Kane, E. E., Poirier, A., Klonoski, K., et al. (2015). A field protocol for the capture and release of callitrichids. Neotropical Primates, 22(2), 59–68.

Watsa, M., Erkenswick, G. A., Pomerantz, A., & Prost, S. (2019, March 11). ddRADSeq in a field setting. Protocols.Io. http://doi.org/10.17504/protocols.io.y2efybe

Wilson, E. O. (1987). The arboreal ant fauna of Peruvian Amazon forests: A first assessment. Biotropica, 19(3), 245. http://doi.org/10.2307/2388342

Zaaijer, S., & Erlich, Y. (2015). Integration of mobile sequencers in an academic classroom, 1–19. http://doi.org/10.1101/035303

Zeng, Y., & Martin, C. H. (2017). Oxford Nanopore sequencing in a research-based undergraduate course. bioRxiv. http://doi.org/10.1101/227439

