## Appendix B for "Genomics in the jungle: using portable sequencing as a teaching tool in field courses"

Course Syllabus


#### Instructors:

##### Gideon Erkenswick

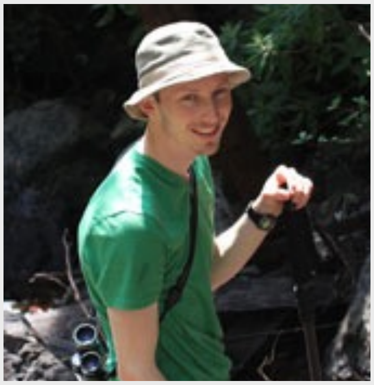

Ph.D., Ecology, Evolution and Systematics, UMSL

##### Mrinalini Watsa

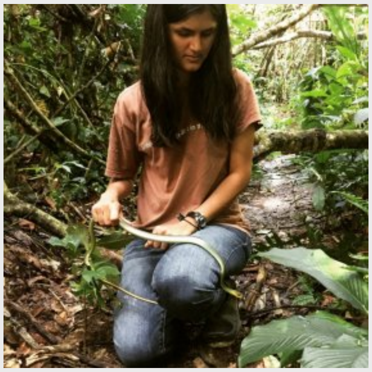

Ph.D., M.A., Biological Anthropology


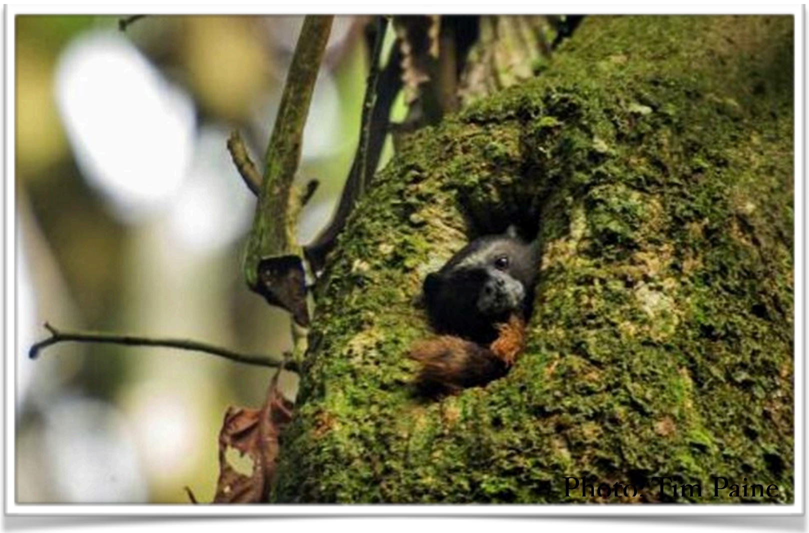


#### Course Introduction

Biological and primatological research is turning to genetic research methods for a deeper look into the biological factors that encode behavior and physiology. We use genetic techniques to determine species delimitations, define populations, understand mating systems, explain behavioral differences in foraging efficiency, screen for disease, conduct paternity studies, evaluate immune status and functioning, and explore microbiome diversity... and these are just a few examples of the full breadth of the field as applied to wildlife biology. The field of genetics is revolutionizing biological research, and in the past few years we have even witnessed the successful deployment of instruments that enable molecular work to be conducted 'on-the-fly' and in the field. These new tools are minimizing the hassles and barriers associated with transporting samples around the world to distant labs that possess the equipment and resources to extract, amplify, and sequence DNA. In many ways, this new technology is democratizing wildlife research by empowering field scientists all

#### Aaron Pomerantz

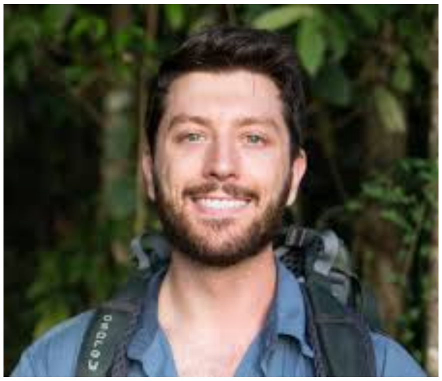

Ph.D. candidate, Evolution

#### Stefan Prost

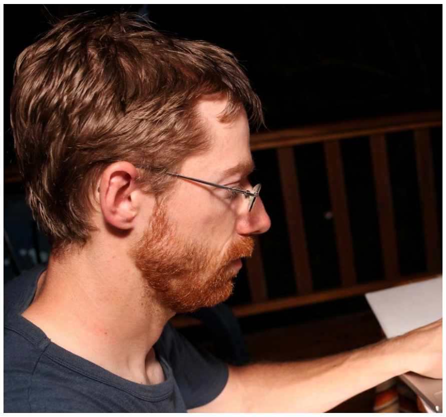

Ph.D., Biology

#### Location:

#### Inkaterra Guides Field Station

#### Course Dates:

July 22 to August 4 , 2018

around the world with genetic tools to directly advance their research and conservation initiatives.

This course will take students to the Peruvian Amazon, where they will learn how field research is conducted, assist in sample collection, and then actually extract, amplify, sequence, and interpret genetic data to answer several practical research questions about wildlife ecology and natural history. It will take place at the Inkaterra Field Guides Station, which is the site of the Green Lab, the world's first tropical rainforest molecular genetics laboratory. Participants will go from sample collection to sequence analysis directly in the rainforest, which is the science of the future. This course is will provide an introduction to next generation primatologists and biologists, who will gain not only the skills requisite for field research but the technical know-how to employ genetic research tools in the field.

### Case Studies

In this course, we will focus on three specific cases (class size and time permitting) in which cutting-edge genomics can help us solve mysteries common to wildlife research in the field. The ultimate goal of all of these projects will be to use a MinION, a USB-sized powerful sequencer that is revolutionizing how we do genomics in some of the craziest places on the planet.

The MinION has been used to:

• do whole-genome sequencing to identify species with PCRs in under 40 minutes and whole genome amplification in under 100 minutes

• conduct real-time DNA sequencing in a remote rainforest by a course instructor

• sequence DNA offline in the Antarctic and even on the International Space Station in outerspace!

##### *Case 1: DNA Fingerprinting*

In 1986, the first criminal was captured on the basis of DNA fingerprinting, a process developed at the University of Leicestershire in the UK. Semen stain samples from two murders were collected and matched to the same perpetrator. An innocent man accused of the crimes was exonerated, and the real culprit caught. Since then, DNA fingerprinting has revolutionized the way crime scenes are processed and biological samples screened. But DNA fingerprinting is not restricted to forensic science applications alone. You can fingerprint anything, and one way in which we do so in wildlife biology is to use a barcode database.

For this case study, we will collect fecal samples from wild primates and use these known samples to identify a series of unknown samples down to the species. To do this, we will barcode the DNA of each sample, and compare it to a reference database. A second dataset will use a similar process to identify unknown plants, keeping in mind that plants are polyploids and so a bit more complex to barcode.

Skillsets: Sample collection from wild primates and Amazonian plants, sample storage and sterile technique, DNA extraction using multiple kits for comparison, amplification of barcoded regions in the genome, gel electrophoresis to confirm amplicon sizes, sequencing of the regions using a MinION, real-time species identification with the WIMP workflow.

#### *Case 2: Does captivity alter primate microbiomes?*

A microbiome is a collection of the genes of all the microbes in a community, and a disturbance of the "normal" microbiome of the gut of an animal (including humans) can result in disease. However, what is normal? Which microbes are present and what do they do? Although much of this information is now known for humans, it is completely unknown in wild primates.

For this case study, we will compare fecal samples collected from a local wildlife rehabilitation center (including animals thought to be healthy and sick) with samples from wild primates living in forests nearby. The goal is to conduct 16s sequencing of all of the bacteria present in these samples. We will also visit the Taricaya Rehabilitation Center to discuss wildlife rehabilitation and the wildlife trade in Peru with veterinarians and keepers on site.

Skillsets: Sample collection (wild and captive), storage and sterile technique, DNA extraction, PCR amplification of the genetic material in the 16S ribosomal subunit, multiplexing of samples and cDNA library prep for sequencing on the MinION, subsequent analysis of data using the WIMP workflow.

#### *Case 3: Environmental DNA*

Imagine, if you will, the contents of a primate fecal sample. Beyond thinking about the icky side of things, a fecal sample is as valuable as gold to a research scientist. It contains a multitude of organisms, each with their own unique trace of DNA. For example, represented will be the bacterial microbiome of the animal's gut, the parasites it might contain, the plant and animal material it consumed, and it's own gut cells with the host's DNA. So from one single sample, we can not only identify the producer of the sample but also everything that animal ate. For a wildlife biologist studying hard-to-see and often unhabituated animals, this is indeed, an unappreciated treasure of information. Even the best animal trackers cannot pursue every animal so closely as to see everything it eats, including tiny insects or rare species of vines in the canopy of a rainforest. Enter, genomics in the jungle!

In this case study, we will use markers such as that of the 18s ribosomal subunit to identify all prokaryotic and eukaryotic DNA in a range of fecal samples. Our goal is to ignore the obvious, such as host DNA but to focus on environmental traces of DNA.

Skillsets: Sample preparation and collection, DNA extraction, library preparation, sequencing on the MinION and analysis.

##### *We will try to answer a few additional questions with all case studies:*

- Can we use the What's In My Pot (WIMP) workflow to accurately classify these species to a reference database in real-time for all of the case studies?
- How many different kinds of samples can we multiplex at one go on a MinION?
- Does the choice of DNA extraction kit affect the outcome?
- Does the length of time we let the MinION run for affect the accuracy of our species identifications?

### Course Objectives

The goals of this course are to give participants advanced training in field techniques important to the collection of biological samples from Amazonian wildlife, their prey and their parasites, all the way to sequencing DNA from these sources.

The course has the following broad objectives:

* To engage in both independent and team-based data collection
* To teach sample collection techniques from Amazonian wildlife, with a focus on primates as model megafauna
* To learn sample storage and clean-lab protocols in the field
* To extract DNA in a field laboratory
* To test DNA quality and quantify it
* To run basic PCRs for a range of markers using multiple protocols on field PCR machines (smaller, lighter, lower-scale and more rugged than typical lab-based machines)
* To explore metagenomics in the field using three case studies:
  * DNA fingerprinting
  * Microbiome metagenomics
  * Environmental DNA

### Course Topics

| Topic of Study | Activity | Description |
|---|---|---|
| **I. Introduction** | | |
| > Threats to the Amazon in the Madre de Dios Department of Peru; conservation efforts of the Amazon Conservation Association (ACA); conservation efforts of FPI | Lecture | A review of the major conservation approaches in the MDD, including the conservation and research efforts of ACA and FPI. |
| > Field ethics, safety precautions, rules,and useful tips. | Discussion | Keeping your footprint to a minimum while working with wildlife in the tropics, and ensuring your safety and that of the wildlife around you. |
| > DNA sequencing and genomics | Lecture/Discussion | An introduction to genomics and its practical applications in the field |
| > Genomics and Ethics | Discussion | What should we be thinking about when applying genomics to wildlife research? Whose rights come into question? Do different countries regulate genomics in different ways? |
| **II. Navigation and Space Use** | | |
| > Basic functions of a handheld GPS and compass | Demonstration | Getting familiar with the most important pieces of equipment you will have in the field. |
| > Waypoint and track data and how to use them | Exercise | Recording key features of the research station with waypoints and tracks |
| > Visualizing spatial data | Exercise | Manipulation of GPS data; creating a digital field map (laptop with USB port required for each participant) |
| **III. Collecting Biological Specimens** | | |
| > Field methodology: indirect observation and biological sampling | Lecture/Practical Exercise | Tracking primates, identifying plants based on botanical features, and collecting non-contaminated samples from each. |
| > Field methodology: sample storage and preparation | Lecture/Practical Exercise | Sample collection methods in the field, time-scales for DNA deterioration, and field laboratory sterile technique |

| Topic of Study | Activity | Description |
|---|---|---|
| **IV. Basic Field Laboratory Techniques** | | |
| > Basic laboratory techniques | Practical Exercise | Regardless of participant background, we will spend a moment practicing good pipetting techniques, sample volume calculations, and gel loading techniques. Repetition is key, getting everyone onto the same skill level |
| > Laboratory Safety | Lecture/Practical Exercise | Even in the field, lab safety is critical. We will go over protocols, cautionary tales, and specific examples of how not to hurt yourself in a field laboratory. |
| > Lab recipes | Practical Exercise | We will learn the processes behind the various lab protocols or recipes we will be following in this course |
| **V. Genomics in the Jungle** | | |
| > DNA barcoding | Lecture | The history of species identification using barcodes, including some of the most exciting applications in wildlife science |
| > Environmental DNA | Lecture | DNA is everywhere, in everything - so why would we care about picking up trace DNA? What are the useful applications of this technique? |
| > Microbiome analyses | Lecture | Microbiome analyses can tell us a great deal about human health - can the same be said for animals? What is a normal monkey gut even look like? |
| **VI. Genomics in a Jungle Lab** | | |
| > DNA extraction | Practical Exercise | Whole genome amplifications and genomic DNA extraction |
| > DNA quantification | Practical Exercise | How much DNA do you have? |
| > PCRs | Practical Exercise | Amplifying markers for all three case studies using PCRs in a field laboratory |
| > Gel electrophoresis | Practical Exercise | Testing if your PCR worked using gel electrophoresis and PCR product quantification |

| Topic of Study | Activity | Description |
|---|---|---|
| > Library prep | Practical Exercise | Creation of DNA libraries from samples |
| > Multiplexing and indexing | Practical Exercise | Minimising sequencing costs by running multiple samples in a single run - multiplexing and indexing samples to tell them apart afterwards |
| > Using the MinION | Practical Exercise | Learn to run sequences on a MinION device - cutting edge genetic sequencing using nanopore flowcells |
| > Basic bioinformatics | Practical Exercise | How to interpret data from the MinION - both real-time and post-hoc |
| **V. Excursions and Activities (time and weather permitting)** | | |
| > Canopy Walkway | Excursion | Participants will be given the opportunity to climb and view the Amazon rainforest from network of rainforest canopy walkways. |
| > Wildlife Hike | Excursion | We will visit Lake Sandoval to look for giant river otters and aquatic birds. |
| > Taricaya Rehabilitation Center | Excursion | The illegal wildlife trade is blooming in Peru, and rehabilitation centers are the final stop for those animals lucky to be recovered - but is being in captivity changing them irrevocably? |
| > Guest Lectures | Lecture | The IGFS is a popular location for field researchers. Participants will be given the opportunity to be exposed to various research groups and interests during their time here, including projects that directly emphasize wildlife biology. |

### Daily Schedule

| Date | Activities | Reading Discussion | Assignments Due |
|---|---|---|---|
| 22nd July | **Arrive in Puerto Maldonado**<br>- Go directly to the IGFS by boat<br>**Night:**<br>- Get to know your peers and instructors | *Discussion:* Field ethics, safety, rules, and tips | |
| 23rd July | **Morning**:<br>- Map the trails: a navigation exercise<br>**Afternoon:**<br>- Welcome to the Green Lab<br>- Lab safety protocols<br>- A lab skills refresher<br>**Night:**<br>*Lectures*<br>- Introduction to the Madre de Dios, ITA & FPI | **Papers**: Swenson et al, 2011; Swamy et al., 2011 | |
| 24th July | **Morning:**<br>- **<u>Group 1</u>**<br>- Sandoval Lake<br>- **<u>Group 2</u>**<br>- Sample storage in the field<br>- Recipes for success<br>- DNA extraction lab 1 (overnight lysing)<br>**Afternoon**:<br>- **<u>Group 1</u>**<br>- Sample storage in the field<br>- Recipes for success<br>- DNA extraction lab 1 (overnight lysing)<br>- **<u>Group 2</u>**<br>- Explore the canopy walkway<br>- Navigating off trail and field sampling<br>**Night**: Lecture<br>- Genetics and wildlife biology | *Discussion*: Lab reports and formats. Group assignments<br>**Paper**: Zeng and Martin, 2017; Erwin 1982 | |

| Date | Activities | Reading Discussion | Assignments Due |
|---|---|---|---|
| 25th July | **Morning:**<br>- **<u>Group 1</u>**<br>- Lab: Complete overnight extractions<br>- **<u>Group 2</u>**<br>- Sandoval Lake<br>**Afternoon**:<br>- **<u>Group 1</u>**<br>- Explore the canopy walkway<br>- Navigating off trail and field sampling<br>- **<u>Group 2</u>**<br>- Lab: Complete overnight extractions<br>**Night**:<br>*Pre-dinner*: Genomics and wildlife biology<br>*Post-dinner*: Introduce the Case Studies. | | Digital Lab Notebook: Due at the end of the night. |
| 26th July | **Morning:**<br>- Visit Taricaya Rehab Center<br>- Sample primates in captivity<br>**Afternoon**:<br>- FTA extractions<br>**Night**:<br>*Lecture:* Conservation Genetics | *Discussion*: The wildlife trade in Peru | Quiz 1: The history of genomics, DNA extraction, navigation techniques, basic lab calculations |
| 27th July | **All Day:**<br>- Split into lab teams<br>- PCR setup and amplifications for various markers<br>- Field activities while PCRs are running<br>- PCR cleanups<br>**Night**<br>*Protocol Troubleshooting and Planning* | **Paper**: Chiou et al., 2018 | |
| 28th July | **All Day:**<br>- Continue labwork from the 27th<br>- Protocol troubleshooting and planning | **Paper**: Thrasher et al., 2018 | Digital Lab Notebook: Due at end of the night. |
| 29th July | **Morning and Afternoon:**<br>- Barcoding PCRs for Multiplexing<br>- PCR cleanups<br>- Normalising Samples<br>- Pooling into Libraries | **Paper**: Cusco et al., 2017 | |

| Date | Activities | Reading Discussion | Assignments Due |
|---|---|---|---|
| 30th July | **Morning and Afternoon:**<br>- Continue labwork from the 30th | Paper: Krehenwinkle et al., 2018; Pomerantz et al., 2018 | Quiz 2: Case studies histories, lab methods: DNA extraction, PCRs, gels |
| 31st July | **Morning and Afternoon:**<br>**Earliest Ready Projects**<br>- Library prep<br>- Trail hikes in between lab<br>- End the day by starting MinION runs<br>**Remaining Projects**<br>- Finish protocols<br>**Night**:<br>- Real-time results with the MinION | | Digital Lab Notebook: Due when library prep is done for your project. |
| 1st August | **Morning and Afternoon:**<br>- Examining real-time sequence results<br>- Exploring MinION outputs<br>- Complete all MinION runs for projects<br>**Night**:<br>- Real-time results with the MinION | | |
| 2nd August | **Morning:**<br>- Data analysis with Case Studies<br>**Afternoon**:<br>- Time to practice presentations and prep for final<br>**Night**:<br>*Final Quiz* | Turn in Draft 1 of your section of the course paper | Final quiz: covers everything learned, focus on the last third of the class |
| 3rd August | **Morning:**<br>- Final excursions and hikes<br>- Completing your projects<br>- Preparing for presentation<br>**Afternoon**:<br>- Time off to pack<br>**Night**:<br>*Final presentations: 10 mins per group* | | |
| 4th August | **Depart from the field station for your flights** | | |

### Course Work

##### *Lab notebook (250 pts)*

Although paper notebooks in a lab are soon becoming a thing of the past, in a field setting with unreliable internet or electricity, it is hard to keep electronic lab notebooks. We will use lab protocols and templates, pre-designed and printed when needed. Participants will maintain a lab notebook per group, as well as individual electronic records of their independent work. At the end of the course, your lab notebook will form a part of your grade, as maintaining an accurate record of the work you do in a lab is a crucial part of being a good research scientist.

##### *Lab Exercises (250 pts):*

Every case study is going to involve a series of lab sessions, each of which will have reports due at the end. These may be recorded in electronic notebooks or turned in by hand, depending on the exercise. Your ability to complete the exercises, remain enthusiastic and a good team player will determine your grade for each lab exercise.

##### *GPS Mapping (100 pts)*

Several activities will involve recording GPS data and manipulating it in the Garmin Basecamp program, which is available for free download. At the end of the course, students will turn in the data from these exercises, which will include tracks made during off-trail hikes and primate follows, as well as a student-generated map of the field station.

##### *Case Study Report and Discussion (200 pts)*

At the end of the course, we will have a roundtable discussion wherein each group will present the outcome of their research in the course. We will discuss the difficulties they faced, their recommendations for repeating this research in the future, and their own interpretation of their success with the project. Each group will also turn in as much of the methods section of a manuscript as they can write within the timeframe, paying particular note to details that will be hard to recover after their departure from the lab itself.

### Reading List (subject to change)


Clayton, J.B., Vangay, P., Huang, H., Ward, T., Hillmann, B.M., Al-Ghalith, G.A., Travis, D.A., Long, H.T., Van Tuan, B., Van Minh, V. and Cabana, F., 2016. Captivity humanizes the primate microbiome. Proceedings of the National Academy of Sciences, p.201521835.

Yildirim, S., Yeoman, C.J., Sipos, M., Torralba, M., Wilson, B.A., Goldberg, T.L., Stumpf, R.M., Leigh, S.R., White, B.A. and Nelson, K.E., 2010. Characterization of the fecal microbiome from non-human wild primates reveals species specific microbial communities. PloS one, 5(11), p.e13963.

Bohmann, K., Evans, A., Gilbert, M.T.P., Carvalho, G.R., Creer, S., Knapp, M., Douglas, W.Y. and De Bruyn, M., 2014. Environmental DNA for wildlife biology and biodiversity monitoring. Trends in Ecology & Evolution, 29(6), pp.358-367.

Mezzasalma, V., Bruni, I., Fontana, D., Galimberti, A., Magoni, C. and Labra, M., 2017. A DNA barcoding approach for identifying species in Amazonian traditional medicine: The case of Piri-Piri. Plant Gene, 9, pp. 1-5.

Janjua, S., Fakhar-I-Abbas, William, K., Malik, I.U. and Mehr, J., 2017. DNA Mini-barcoding for wildlife trade control: a case study on identification of highly processed animal materials. Mitochondrial DNA Part A, 28(4), pp.544-546.

Chiou KL, Bergey CM. 2018. Methylation-based enrichment facilitates low-cost, noninvasive genomic scale sequencing of populations from feces. Scientific Reports 8:1975.

Cusco A, Vines J, D'Andreano S, Riva F, Casellas J, Sanchez A, Francino O. 2017. Using MinION to characterize dog skin microbiota through full-length 16S rRNA gene sequencing approach. biorxiv.

Krehenwinkel H, Pomerantz A, Henderson JB, Kennedy SR, Lim JY, Swamy V, Shoobridge JD, Patel NH, Gillespie RG, Prost S. 2018. Nanopore sequencing of long ribosomal DNA amplicons enables portable and simple biodiversity assessments with high phylogenetic resolution across broad taxonomic scale:1–24.

Pomerantz A, Penafiel N, Arteaga A, Bustamante L, Pichardo F, Coloma LA, Barrio-Amoros CL, Salazar-Valenzuela D, Prost S. 2018. Real-time DNA barcoding in a rainforest using nanopore sequencing: opportunities for rapid biodiversity assessments and local capacity building. GigaScience 7:74–14.

Zeng Y, Martin CH. 2017. Oxford Nanopore sequencing in a research-based undergraduate course. bioRxiv.

### Grading Criteria

Individual and group assignments will be assessed according to the following point schedule:

| Assessment Item | Date Due | Points Possible | Total Points Possible |
| --- | --- | --- | --- |
| GPS map and track log | Students are to continuously maintain a GPS tracks log | 100 | 1000 |
| Lab notebook | The final report | 250 | |
| Lab exercises | Reports from various exercises, benchling or paper | 250 | |
| Quiz 1 | | 100 | |
| Final Exam | | 100 | |
| Case Study Report | | 200 | |
