## Appendix C for "Genomics in the jungle: using portable sequencing as a teaching tool in field courses"

**Appendix C: Post Course Questionnaire (PCQ)**

### Course Evaluation: (Genomics)

Please give us your opinion of the course in the form below. We loved having you participate in the course, and would love to know how it could be improved for future generations of students. Thank you!

**Please enter your course name below** *

**Please enter your course month and year below** *

Eg: January, 2018

### Section 1 of 4: Do you agree with the following statements?

**The readings helped me learn the course material** *

- Strongly disagree
- Disagree
- Neutral
- Agree
- Strongly Agree
- Not Applicable

**The lectures helped me learn the course material** *

- Strongly disagree
- Disagree
- Neutral

- ◯ Agree
- ◯ Strongly Agree

**The material covered in the course was challenging to me** *

- ◯ Strongly disagree
- ◯ Disagree
- ◯ Neutral
- ◯ Agree
- ◯ Strongly Agree

**The course activities were challenging to me** *

- ◯ Strongly disagree
- ◯ Disagree
- ◯ Neutral
- ◯ Agree
- ◯ Strongly Agree

**I received enough attention from the instructor(s)** *

- ◯ Strongly disagree
- ◯ Disagree
- ◯ Neutral
- ◯ Agree
- ◯ Strongly Agree

**The course instructor(s) were sufficiently knowledgeable about the topics taught** *

- ◯ Strongly disagree
- ◯ Disagree
- ◯ Neutral
- ◯ Agree

- ○ Strongly Agree

**Overall, I am satisfied at the end of this course in terms of my expectations when I signed up for it** *

- ○ Strongly disagree
- ○ Disagree
- ○ Neutral
- ○ Agree
- ○ Strongly Agree

**This course has helped me figure out more about my interests and possible future goals** *

- ○ Strongly disagree
- ○ Disagree
- ○ Neutral
- ○ Agree
- ○ Strongly Agree

**I would recommend this course to someone else** *

- ○ Strongly disagree
- ○ Disagree
- ○ Neutral
- ○ Agree
- ○ Strongly Agree

### Section 2 of 4: For the following questions, please select ALL that apply:

---

**When would be the right time to take this course?** *

- ☐ High school graduate (age 17)
- ☐ Freshman in College (Year 1, age 18)

- [ ] Sophomore in College (Year 2, age 19)
- [ ] Junior in College (Year 3, age 20)
- [ ] Senior in College (Year 4, age 21)
- [ ] College graduate (age 22+)
- [ ] Graduate student
- [ ] Professional Advancement
- [ ] No Opinion

**Please pick as many of these words that apply to the experience you had on this course:** *

- [ ] Amazing
- [ ] Regret
- [ ] Useless
- [ ] Fun
- [ ] Tough
- [ ] Satisfied
- [ ] Fascinating
- [ ] Difficult
- [ ] Unforgettable
- [ ] Boring
- [ ] Useful
- [ ] Forgettable
- [ ] Valuable
- [ ] Tedious
- [ ] Dissatisfied

**Please check if your instructor(s) did any of the following:** *

- [ ] discussed the FPI sexual harassment policy with participants as a group
- [ ] discussed the FPI sexual harassment policy with you individua

- [ ] discussed field safety protocols with the group
- [ ] discussed wildlife handling ethics with the group
- [ ] listened and reacted to concerns you brought up regarding course content
- [ ] listened and reacted to concerns you brought up regarding course social dynamics
- [ ] behaved ethically regarding wildlife handling on the course
- [ ] None of the above

Check as many as apply

### Section 3 of 4: Please respond to the following questions, and feel free to be as concise or elaborate as you like.

**How would you rate your level of experience with molecular biology techniques before taking this course?**

**What challenges (if any) did you encounter with yourself, or in the group, while performing experiments in the lab?**

**In your opinion, how clear were the scientific objectives and methods? If you were to do this all over again, what might you change to make things more clear or straightforward?**

**Now that you have completed the two week "Genomics in the Jungle" course, do you feel more proficient with any of the topics or experiments that you completed? If so, which ones, and which areas do you still wish to work on further?**

**Real-time portable DNA sequencing can have many applications. What would you consider to be the top 3 most important implementations with portable scientific instruments? Are there any tools that you think could be made more cost-effective / portable in the future?**

### Section 4 of 4: Please comment on the following and let us know how we did, and if we can improve. Your opinion is very important to us.

**The facilities at each of the field sites used on this course.** *

**Please list at least 2 (more are welcome) specific course activities/events/experiences you enjoyed the MOST and the LEAST.** *

Eg: Most enjoyable:.... Least enjoyable:....

**Please share any specific changes/modifications/additions/eliminations to improve the course itinerary/schedule/activities/exercises.** *

**Please indicate both POSITIVE and NEGATIVE qualities of the course instructor(s) that could be retained/improved on in future courses.** *

**Please list the most useful/interesting skills or knowledge you gained during the course** *

**The cost effectiveness of the course (in comparison to other similar opportunities)** *

**Please note your most vivid memories from the course (positive or negative)** *

**Any other comments?**

Specifically, were there other words you would use to describe your experience on this course?

**Would you be interested in contributing a short article for the FPI blog on any of your experiences while attending this course? ***

- Yes
- No
- Not sure, tell me more
