## Appendix A for "Genomics in the jungle: using portable sequencing as a teaching tool in field courses"

| Item | Number |
|---|---|
| Basic Infrastructure | |
| Autoclave | 1 |
| Oven | 1 |
| Microwave | 1 |
| Lab Computer | 1 |
| Refrigerator with small freezer | 1 |
| Freezer | 1 |
| Steam Water Distiller - 1 Gallon | 1 |
| AmScope 100X-1000X Trinocular LED Infinity Plan Phase Contrast Microscope | 1 |
| UV Light | 1 |
| Voltage Converter | 1 |
| Benchtop Equipment | |
| Vortexer | 2 |
| pH Meter | 1 |
| Mini Microcentrifuge | 2 |
| Magnetic bead separation rack 1.5ml tubes | 1 |
| Magnetic bead separation rack 96-well plate | 1 |
| Dry Bath Incubator | 1 |
| Water Bath | 1 |
| 8-well PCR machine | 5 |
| 16-well PCR machine | 2 |
| Bluegel Electrophoresis | 3 |
| MiniOne Electrophoresis Package | 2 |
| Quantus Fluorometer | 1 |
| Balance | 1 |
| Microcentrifuge (24 well) | 2 |
| Cool Cube | 1 |

| Item | Number |
|---|---|
| Vaccuum Pump | 1 |
| Large Centrifuge (Temp controlled) | 1 |
| MinION Sequencer | 1 |
| Pipettes | |
| Micropipette (20-200uL) | 6 |
| Micropipette (100-1000uL) | 6 |
| Micropipette (1-10uL) | 6 |
| Multichannell pipette 20-200ul | 2 |
| Pipette pumps (2ml, 5ml, 10ml) | 1 set |
| Course-Specific Ingredients | |
| Taq polymerase (dNTPs and Magnesium included) | 2 |
| PCR master mix | 1 |
| Quantus Fluorometer Assays | 1 |
| PCR strip tubes | 250 |
| 1.5mL Microcentrifuge Tubes | 1000 |
| 96 Barcoding Kit | 1 |
| Gelgreen | 1 |
| Gel loading dye | 1 |
| Agarose | 1 |
| TBE Buffer 10X | 1 |
| Gel extraction kit | 1 |
| Cutsmart Buffer | 1 |
| Restriction enzyme | 1 |
| ATP | 1 |
| T4 DNA Ligase | 1 |
